# Gene Set Analysis for time-to-event outcome new approach based on the Generalized Berk–Jones statistic and comparison with existing methods

**DOI:** 10.1101/2021.09.07.459329

**Authors:** Thomas Ferté, Laura Villain, Rodolphe Thiébaut, Boris P. Hejblum

## Abstract

Gene set analysis evaluates the collective impact of groups of genes on an outcome of interest, such as disease occurrence. By incorporating biological knowledge through predefined gene sets, this approach enhances the interpretability of results and improves statistical power compared to gene-wise analyses. In the context of time-to-event data, existing methods are limited and fail to account for potentially strong correlations within gene sets. Given the strong performance of the Generalized Berk-Jones (`GBJ`) statistic, which effectively incorporates correlation within the test statistic, we adapted this method to the time-to-event framework using a Cox model. We then compared its performance with established methods, including the `Wald test`, `global test`, and `global boost test`.

Our proposed method, `sGBJ`, shows an over-control of Type I error, leading to reduced statistical power compared to other methods in numerical studies. We further benchmarked these methods in two different real-world contexts: gliomas and breast cancer. The Wald test emerged as the most effective, identifying the largest number of significant pathways while maintaining appropriate control of Type I error in simulation settings. `sGBJ` closely followed demonstrating good performances, without a significant loss of statistical power in analyzing these two real-world biomedical datasets.

## 1. Introduction

The analysis of the whole transcriptome thanks to RNA-seq (Wang et al., 2009) is usually based on differential expression methods. Statistical methods such as `edgeR` (Robinson et al., 2010), `DESeq2` (Love et al., 2014), `limma-voom` (Law et al., 2014), or `dearseq` (Gauthier et al., 2020) allow comparing gene expression between groups of patients or within patients over time. However, gene-wise analysis exhibits several limitations: i) the high-dimensionality of gene expression data (often tens of thousands of genes are measured per sample) might lead to either no gene being detected differentially expressed after multiple testing correction, or on the contrary to a very large list of genes that is difficult to interpret biologically; ii) many genes interact within biological pathways, but with potentially only small individual changes affecting a few genes, genes within a pathway can fail to be detected as differentially expressed. To address these issues, Gene Set Analysis leverages predefined sets of genes – available in databases such as MSigDB (Liberzon et al., 2011), KEGG (Ogata et al., 1999) or Gene Ontology (Ashburner et al., 2000) for instance – and identifies groups of genes differentially expressed rather than single genes. The reduction of the number of statistical tests as well as the strength of a coordinated signal within a gene set both increase statistical power, and avoid missing important biological links with the outcome of interest. Numerous methods have been proposed to analyze gene set expression in various context (Maciejewski, 2014), such as Gene Set Enrichment Analysis (`GSEA`) (Subramanian et al., 2005), `GSA` (Efron and Tibshirani, 2007), Time-course Gene Set Analysis (`TcGSA`) (Hejblum et al., 2015), Generalized Higher Criticism (`GHC`) (Barnett et al., 2017), `dearseq` (Agniel and Hejblum, 2017) or the Generalized Berk-Jones (`GBJ`) statistic (Gaynor et al., 2019).

In the time-to-event context, some methods are able to tackle the high-dimensionality of gene expression data, such as survival random forest (Ishwaran et al., 2008) or penalized Cox regression (Tibshirani, 1997), but only a handful can perform gene set analysis. Based on Kernel Machine, Cai et al. (2011) proposed a score test for survival gene set analysis, that Neykov et al. (2018) later adapted to take into account competing risk. Besides, three additional methods are building on the Cox model to perform survival gene set analysis, using different test statistics and relying on permutations to compute the associated p-values: i) the `global test` (Goeman et al., 2005), ii) the `Wald test` (Adewale et al., 2008), and iii) the `global boost test` (Boulesteix and Hothorn, 2010). In their review, Lee et al. (2011) found all three tests outperformed `GSEA` in terms of statistical power; even though the `global test` rely on the strong hypothesis that all regression coefficients are sampled from one unique probability distribution, and none of the three compared tests takes into account the correlation structures between the genes in the pathway.

Gaynor et al. (2019) compared the `GBJ` statistic with `GSA`, `GSEA`, and `GHC` to identify gene sets differentially expressed between two tumor grade, and they concluded that their proposed `GBJ` method was more consistent. The `GBJ` statistic has the advantage of not requiring any distributional assumption on the count data or the regression coefficients, while also taking into account the correlation structure between the genes within the same set. We propose to adapt this `GBJ` statistic in the time-to-event context using a statistic derived from the Cox model. We denote our method by `sGBJ` for “survival Generalized Berk–Jones”.

In this paper, we first describe our new method `sGBJ` in section 2 before benchmarking existing methods together with `sGBJ` in a simulation study inspired from both Lee et al. (2011) and real-world data. We also compare the results of the different methods in two real-world data analyses, with brain cancer data and breast cancer data respectively section 4, and finally discuss our results and their limits in section 5. The `sGBJ` method is available as an `R` package from CRAN at https://CRAN.R-project.org/package=sGBJ.

## 2. Method

### 2.1. The Generalized Berk–Jones set-based testing method

The `GBJ` statistic can be used in a set based testing procedure to determine the association between gene expression in a gene set and a given clinical outcome (e.g: tumor grade). Introduced by Sun and Lin (2017), it was used in the context of Genome-Wide Association Studies (GWAS) (Sun et al., 2019) and its consistency was evaluated for identifying pathways whose expression is associated with either low or high grade of breast cancer (Gaynor et al., 2019). Below is a 4 steps description of this `GBJ` testing procedure:

1. Model and null hypothesis specification for a given gene set of *d* genes: Sun and Lin (2017) study the association between the outcome of interest *Y*_*i*_ of patient *i* with the vector **G**_*i*_ the expression of *d* genes with a generalized linear model, with *µ*_**i**_ the conditional mean of *Y*_*i*_ and **X**_*i*_ the other covariates: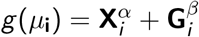.
2. For each gene *j* of a given gene set with *j* = {1, …, *d*} , a score value *Z*_*j*_ is computed. It must verify **Z** ∼ 𝒩 (**0, Σ**) under the null hypothesis.
3. The `GBJ` statistic is computed for the whole gene set based on the *Z*_*j*_ values of each gene, using the threshold function *S* computing the number of genes in the gene set that have an absolute value of their score *Z*_*j*_ higher than a limit *t* and thus defined as 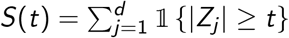.
4. Once the `GBJ` statistic is computed, its associated p-values is determined with a root-finding algorithm that identifies the boundaries points *b*_*j*_ , i.e. the limit value of the *j*^*th*^ greatest absolute value of **Z**.

### 2.2. Extension to time-to-event data

The `GBJ` method as defined by Sun and Lin (2017) can be applied to any generalized linear model, but not for time-to-event data analysis. From the third step onward, the procedure relies on a score statistic calculated for each gene within the gene set, with a multivariate normal distribution assumption on this score centered around zero under the null hypothesis (i.e. no association between gene expression and the dependent time-to-event variable **Y**). We propose a survival `GBJ` method, named `sGBJ`, which uses a score suited to time-to-event in order to compute a `GBJ` statistic in this context.

#### 2.2.1. Model

To deal with a time-to-event outcome, we rely on a Cox proportional hazards regression model:

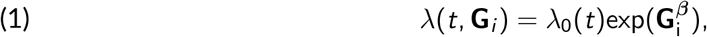

with **G**_*i*_ the vector of gene expressions of patient *i* . The null hypothesis is that there is no association between patient survival and the expression of genes within the set: *H*_0_ : ***β*** = **0**_*d*×1_.

#### 2.2.2. Z score

For each gene *j* of the gene set, we compute the value *Z*_*j*_ as the square root of the Wald statistic (Engle, 1984):

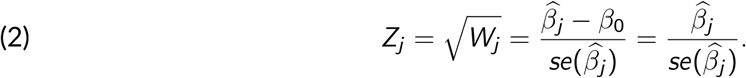

This yields a vector **Z** that follows 𝒩(**0, Σ**) under the null hypothesis *H*_0_. **Σ** is estimated through permutations of the original survival observations: for each permutation *p* ∈ {1, … , *P*}, the 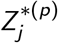 value for each gene *j* gives us a **Z**^∗(*p*)^ vector of length *d* , allowing us to estimate Σ with the empirical covariance across permutations 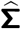.

#### 2.2.3. GBJ statistic

The `GBJ` statistic is computed according to Sun and Lin (2017):

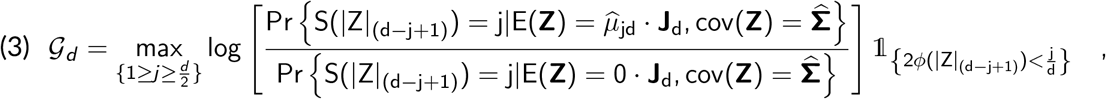

with 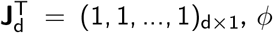 the survival function of a standard normal random variable, and 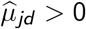 solving the following equation:

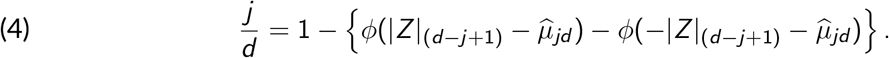

Of note, the `GBJ` statistic 𝒢_*d*_ is calculated only on the higher half values of |**Z**| , and can be represented as the maximum of a set of likelihood ratios on *S* (*t*) (Sun et al., 2019).

#### 2.2.4. p-value computation

The p-value can be computed as:

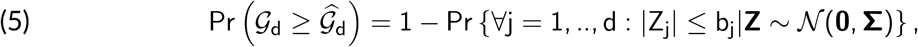

with 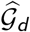 the observed value of the `GBJ` statistic. The associated rejection region is then:

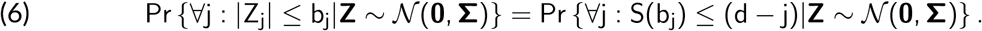

## 3. Numerical studies and comparisons

### 3.1. Numerical simulation scenarios

`sGBJ` does not require any distributional assumption on the gene expression matrix, thus we simulated this matrix for *N* ∈ {50, 100} observations following a multivariate normal distribution 𝒩 (**0, C**), with **C** being the gene covariance matrix. Similarly to Lee et al. (2011), we simulated a gene set of *NG* = {10, 50} genes, among which 20% were significantly associated with survival. We set the variance *C*_*jj*_ = 0.2 for each gene *j*. We generated three different correlations scenarios designed to mimic observed features of real-world data inpired by both Rembrandt (Madhavan et al., 2011) and a breast cancer study (Van De Vijver et al., 2002) datasets:

- **Case (I)**: overall correlation follows a non-standard beta

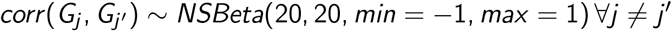
- **Case (II)**: this scenario mimics both Rembrandt and Breats Cancer data (see section 4.1 and 4.2).
  - correlation between significant genes follows a *NSBeta*(10, 10, *min* = −1, *max* = 1),
  - correlation with non-significant genes follows a *NSBeta*(25, 25, *min* = −1, *max* = 1)
- **Case (III)**:
  - correlation between significant genes is 0.2,
  - correlation with non significant genes is 0.

Note that Case (I) and (II) are not guaranteed to generate positive definite matrix. So once the correlation values are sampled, if the matrix is not positive definite, the nearest positive definite matrix is computed using the algorithm defined in Higham (2002).

Survival times were then generated using a Cox model featuring an effect *β* for the significant genes. We investigated three different effect types:

- **Type (A)**: *β*_*j*_ ∼ 𝒩(0, 0.4^2^).
- **Type (B)**: half of the gene effects follow *β*_*j*_ ∼ 𝒩(−0.4, 0.2^2^), while the other half follow *β*_*j*_ ∼ 𝒩(0.4, 0.2^2^). This setting mimics the Rembrandt dataset (see section 4.1).
- **Type (C)**: half of the gene effects follow *β*_*j*_ ∼ 𝒩(−0.8, 0.4^2^), while the other half follow *β*_*j*_ ∼ 𝒩(0.8, 0.4^2^). This setting mimics breast cancer data (see section 4.2).
- **Type (Z)**: ***β*** = **0**, evaluates the type-I error

In all scenarios, the correlation matrix of ***β*** is the same as the correlation matrix of **G**. We also considered potential censoring: censoring times were generated following an exponential distribution for 30% of the observations.

For each scenario, 1, 000 independent Monte-Carlo repetitions were generated.

### 3.2. Compared methods

`sGBJ` was compared to three state-of-the-art methods: i) the `global test` (Goeman et al., 2005), ii) the `Wald test` (Adewale et al., 2008), and iii) the `global boost test` (Boulesteix and Hothorn, 2010), that are also based on statistics derived from the Cox model. The `global test` relies on the hypothesis that all *β*_*j*_ are sampled from the same normal distribution centered in 0 with a common variance *σ*^2^, so the null hypothesis can be reduced to testing *σ*^2^ = 0. The `Wald test` uses the sum of squares of the Wald statistics from a Cox model applied individually to each gene within a gene set, possibly adjusted on other covariates. The `global boost test` combines the Cox model with a boosting algorithm to assess the additional predictive value of gene expression within a gene set. Each method relies on a different test statistic, and the corresponding p-value is computed through permutations for all three methods.

### 3.3. Results

Figure 1 highlights the performance of the four methods across the different simulation settings. Panel A reveals that statistical power varied significantly depending on the scenario, with lower power observed as the number of individuals or genes decreased. The `global test` showed poorer performance in Type C, likely due to the violation of its assumption of normally distributed coefficients. Similarly, `sGBJ` underperformed when the number of genes was high, which may be attributed to its overly conservative control of Type I error.

**Figure 1.**
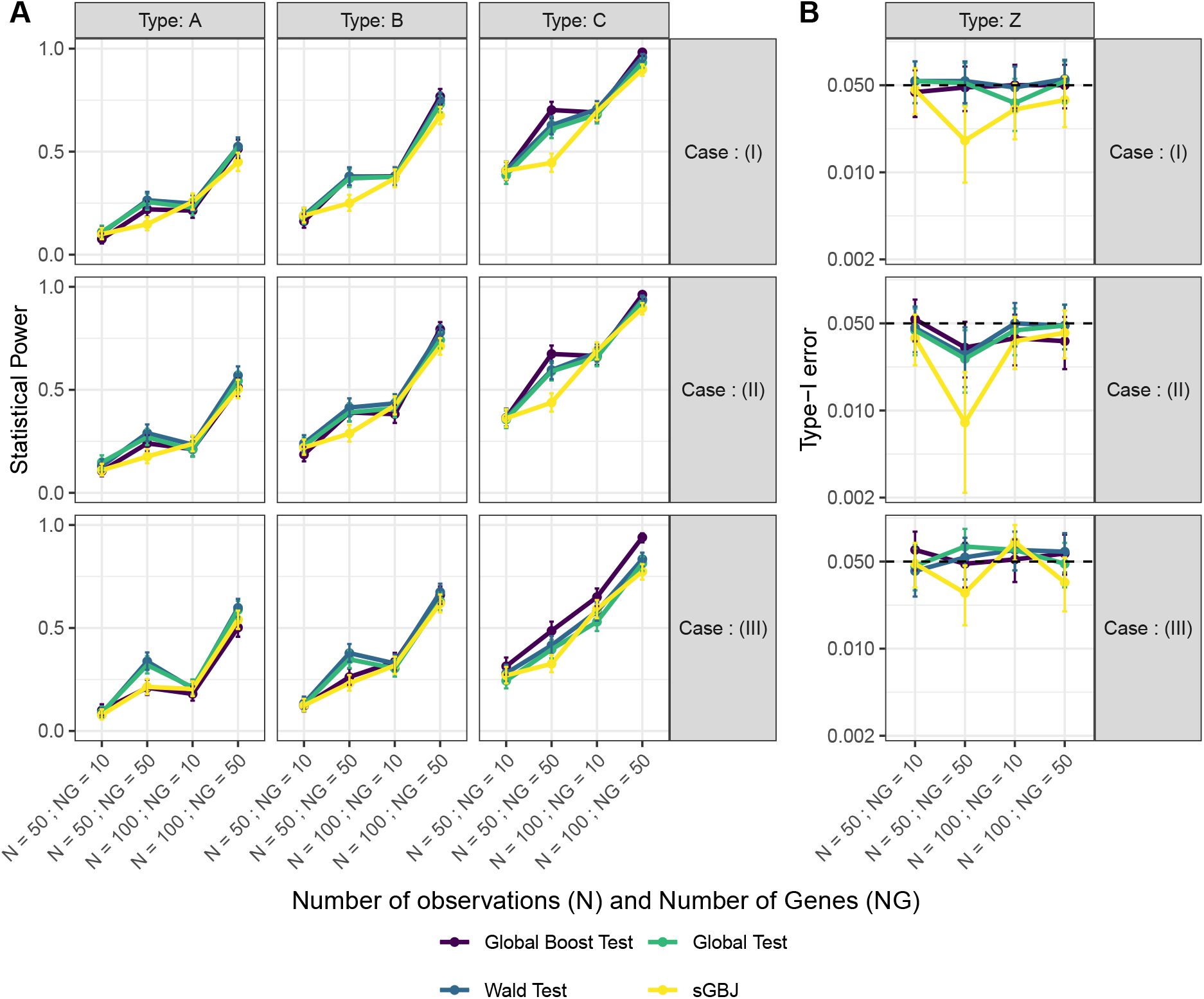
Panel A shows the statistical power of the four methods (`sGBJ`, `Global Boost Test`, `Wald Test`, and `Global Test`) across nine different scenarios, which combine three correlation cases (I, II, and III) with three types of effects (A, B, and C). For more details on these scenarios, refer to Section 3.1. Panel B presents the type I error rates of the methods, stratified by the different correlation cases.

In contrast, the `Wald test` and `global boost test` consistently ranked among the top-performing methods. Although the `global boost test` exhibited weaker performance in Case III--Type A, it excelled in Case III-Type C. This discrepancy may be due to differences in modeling approaches: the `Wald test` fits a separate Cox model for each feature without accounting for correlation structure, whereas the `global boost test` simultaneously models the effects of all genes within the boosting procedure.

Notably, under conditions most relevant for the application (N=100), the methods demonstrated comparable power, with `sGBJ` being slightly more conservative due to its stringent control of Type I error (see Panel B of Figure 1).

## 4. Real-world medical data applications

### 4.1. Survival analysis in glioma subtypes

Glioma represent over 80% of malignant brain tumor, and are associated with poor survival with less than 5% of survival at 5 years for the glioblastoma (the most common type of glioma) (Ostrom et al., 2014). Glioma can be classified in different types according to the World Health Organization (WHO), with mostly histological and molecular alterations criteria (a first classification from 2007 was revised in 2016). The severity can increase from grade I to grade IV, grade IV being mostly the glioblastoma (Chen et al., 2017). The high genetic heterogeneity of glioma, and the poor response to targeted treatment with a high rate of recurrence (Norden and Wen, 2006; Sathornsumetee et al., 2007), makes it hard to treat patients with glioma. To better understand the mechanisms underlying glioma, researchers are actively investigating the molecular and genetic processes linked with the different types of gliomas and patient survival or risk of recurrence.

Rembrandt is one of the largest public datasets on glioma, featuring clinical and genomic data on 671 patients with any types of gliomas (based on the 2007 WHO classification) collected across 14 institutions. This bioinformatic platform for brain cancer research is accessible from the Georgetown University’s G-DOC System and described in *Georgetown Database of Cancer (G-DOC) platform* (n.d.) and Madhavan et al. (2011, 2009). For our application, we restricted our comparison to patients with either one of the two main glioma types: i) astrocytoma, and ii) oligodendroglioma. We included all such patients from Rembrandt, for whom both tumor gene expression (as measured by micro-array) and overall survival follow-up was available. Of note, Glioblastoma patients were excluded to avoid too large genetic variation across compared glioma types. Table 1 presents the different characteristics of the total 154 patients included, and Panel A of Figure 2 displays the corresponding Kaplan-Meier curves stratified on the glioma type. We observe that astrocytoma and oligodendroglioma patients present similar survival curves.

**Table 1.** Characteristics of Astrocytoma and Oligodendroglioma patients included from the Rembrant dataset.

|  | Astrocytoma (108) | Oligodendroglioma (46) |
| --- | --- | --- |
| Age < 40 yo n (%) | 52 (48 %) | 22 (48 %) |
| Male n (%) | 71 (66 %) | 23 (50%) |
| Death n (%) | 85 (79 %) | 34 (74 %) |
| Median follow-up period in months Median (IQR) | 37 (15 ; 66) | 32 (16 ; 48) |

**Figure 2.**
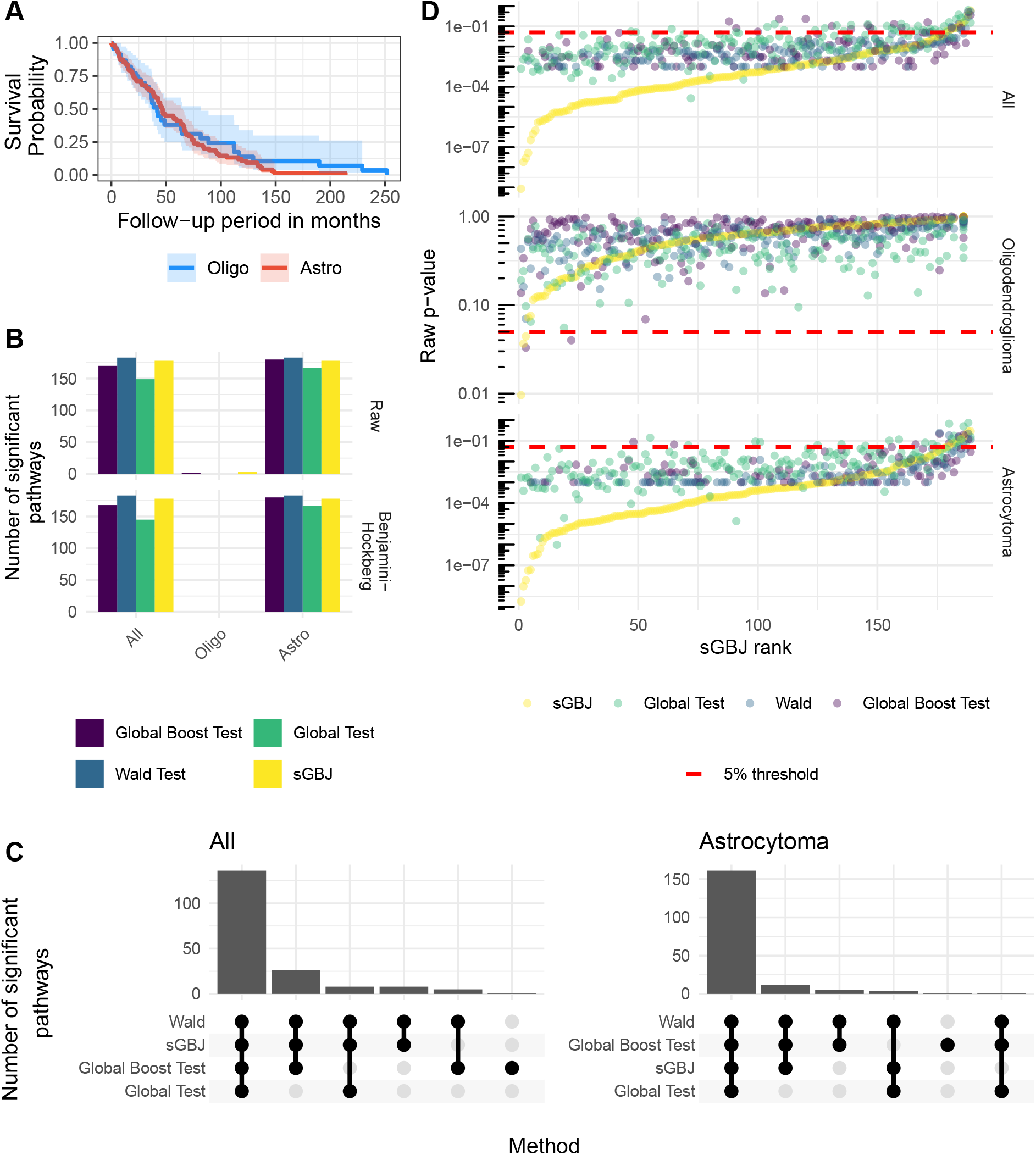
Panel A shows Kaplan-Meier curves for Rembrandt patients, stratified by glioma type. Panel B displays the number of significant pathways identified by the four methods (`sGBJ`, `global boost test`, `Wald test`, and `global test`) for astrocytoma, oligodendroglioma, and the combined cohort with all patients, at the 5% threshold using either raw p-values or adjusted p-values for multiple testing with the Benjmaini-Hochberg procedure. Panel C illustrates the agreement between the significant pathways identified by the different methods. Panel D plots the raw p-values against the ordered ranks of `sGBJ` for all four methods, with a 5% significance threshold. P-values equal to zero were excluded for clarity on the log scale.

We studied the association between patient survival and pathway gene expression, both overall and specifically for the two types of glioma: astrocytoma and oligodendroglioma. We investigated the C6 collection, from MSigDB (Liberzon et al., 2011). It contains 189 gene sets related to cellular pathways that are often found to be disregulated in oncogenic studies. We used the following hazard equation to link the recurrence time with gene expression within a gene set of interest:

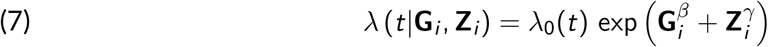

with *λ*_0_(*t*) the baseline hazard function, *G*_*ij*_ the gene expression of the gene *j* for the patient *i* , *β*_*j*_ the associated effect, and *Z*_*il*_ the values of the covariate *l* (i.e. age or sex) with *γ*_*l*_ the associated effect. The null hypothesis tested by the method is that all *β*_*j*_ are zeros within a gene set.

No pathways were identified as significantly associated with survival in oligodendroglioma patients (after multiple testing correction). However, for astrocytoma patients, the `Wald test` identified 96.8% of pathways as significant after applying the Benjamini-Hochberg correction for multiple testing (Benjamini and Hochberg, 1995). Most methods yielded similar results, except for the `global test`, which identified only 76.7% of significant pathways compared to 88.9% for the `global boost test`, 94.2% for `sGBJ`, and 96.8% for the `Wald test`, as shown in Panel B of Figure 2. The methods demonstrated good agreement in identifying significant pathways, as illustrated in Panel C of Figure 2.

Panel D of Figure 2 presents the p-values computed by the different methods. Both the `sGBJ` and `global test` methods use asymptotic p-values (note that the *GlobalTest* `R` package does not support permutation p-values when using a Cox model with covariates). As a result, these methods have greater accuracy compared to those relying on 1,000 permutations, where the lowest non-zero p-value achievable is 1/1,001. While this limitation in permutation number does not significantly affect oligodendroglioma patients (as their p-values are well above 10^−3^), it has a minor impact on astrocytoma patients. This limitation underscores the importance of precise p-value estimation, particularly in multiple testing contexts (Phipson and Smyth, 2010).

### 4.2. Progression free survival in breast cancer

Breast cancer is the most common cancer, with a number of global deaths predicted to reach 11 million per year by 2030 (Benson and Jatoi, 2012). Survival is highly impacted by the presence of metastasis (Scully et al., 2012), which justifies the search for genes or pathways associated with either death or metastasis.

We analyzed gene expression and clinical data from the Netherlands Cancer Institute. Primary breast carcinomas data were available in the `R` package *breastCancerNKI* on bioconductor (Markus Schroeder, 2017), originally reported by Van De Vijver et al. (2002) and later reanalyzed by several teams (Cai et al., 2011; Neykov et al., 2018). This data set comprises 24,481 gene expressions measured by Agilent technology and 337 samples. We removed 42 patients with missing phenotype data. Following Gao et al. (2016), we removed genes with more than 10% missing data and imputed the remaining missing data using nearest neighbor averaging (Trevor Hastie, 2017). Gene expression was further standardized across samples through quantile normalization (as implemented in the *limma* bioconductor package) (Bolstad et al., 2003). The median follow-up period for the 295 remaining patients was 7.2 years, and two events of interest were recorded: the appearance of metastasis and death. We studied metastasis-free survival, i.e., a composite outcome considering time to either metastasis or death, and its potential association with specific gene sets or pathways.

Table 2 presents the different characteristics of the patients included, and Panel A of Figure 3 displays the corresponding Kaplan-Meier curves.

**Table 2.** Characteristics of the 295 patients included in our metastasis free survival analysis of data from Van De Vijver et al. (2002)

|  | Overall (260) |
| --- | --- |
| Age in years Median (IQR) | 44 (40 ; 49) |
| Grade |  |
| 1 | 75 (25 %) |
| 2 | 101 (34 %) |
| 3 | 119 (40 %) |
| Event - metastasis or death n (%) | 106 (36 %) |
| Median follow-up period in years Median (IQR) | 7.2 (5.3 ; 10.3) |

**Figure 3.**
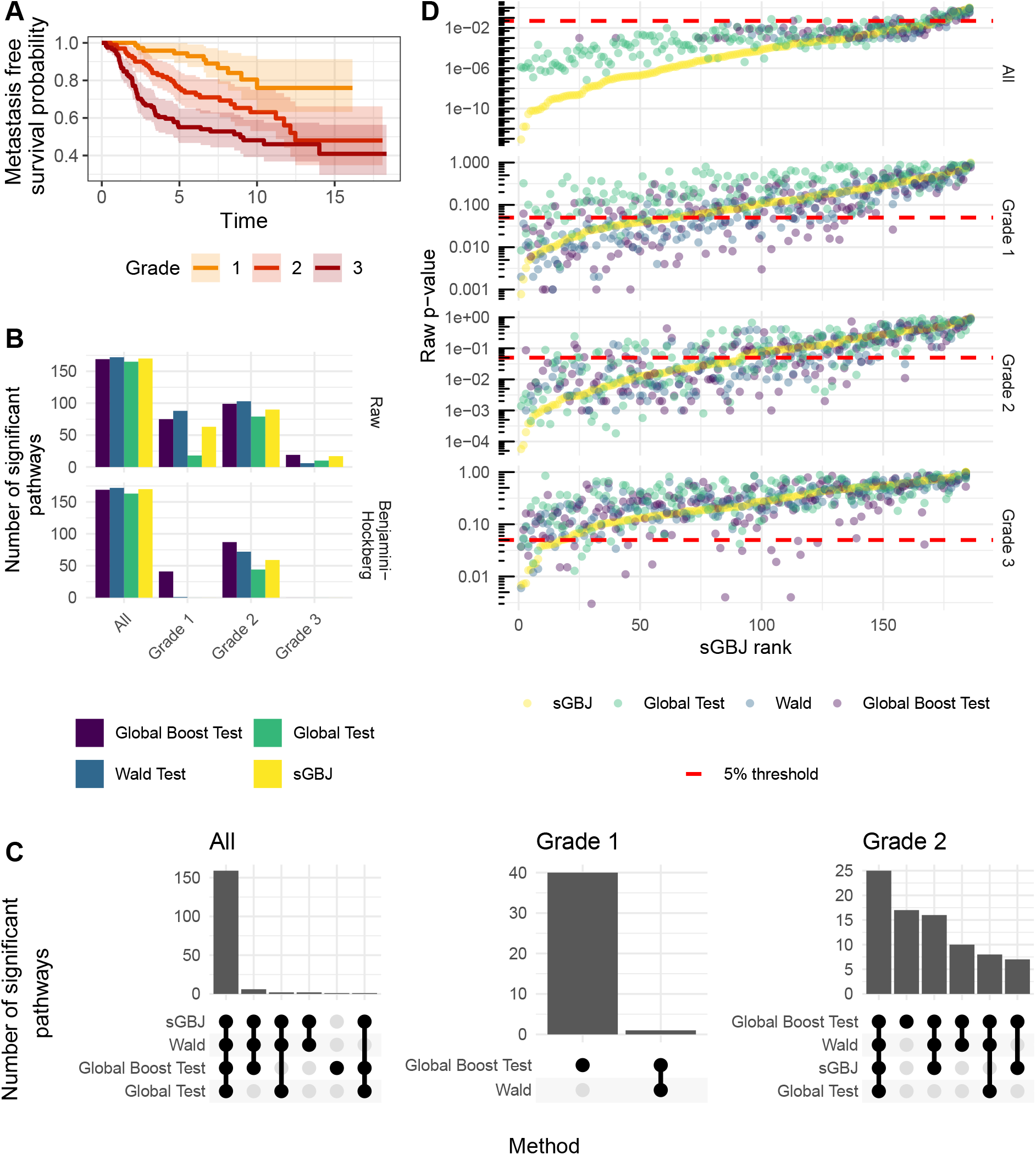
Panel A shows Kaplan-Meier curves for breast cancer patients, stratified by cancer grade. Panel B displays the number of significant pathways identified by the four methods (`sGBJ`, `Global Boost Test`, `Wald Test`, and `Global Test`) for each grade, and the combined cohort with all patients, at the 5% threshold using either raw p-values or adjusted p-values for multiple testing with the Benjmaini-Hochberg procedure. Panel C illustrates the agreement between the significant pathways identified by the different methods. Panel D plots the raw p-values against the ordered ranks of `sGBJ` for all four methods, with a 5% significance threshold. P-values equal to zero were excluded for clarity on the log scale.

We studied the association between the metastasis-free survival and gene set expression adjusted for age (McPherson et al., 2000), with and without stratifying on severity grade using the following Cox model:

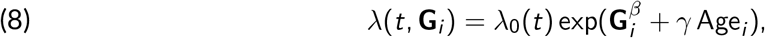

where *λ*_0_(*t*) is the baseline hazard function, *G*_*ij*_ represents the gene expression of gene *j* for patient *i* and *β*_*j*_ is the associated effect. The null hypothesis tested by the method assumes that all *β* values are null.

We investigated 186 pathways from the KEGG database (Kanehisa et al., 2007) corresponding to the version v2.5 of the “Canonical Pathway” (CP) collection from the C2 (“curated gene sets”) category of mSigDB (Molecular Signature Database, Dolgalev (2025). Among these pathways, 92.5%, 91.4%, 90.9% and 87.6% were identified as significant after applying the Benjamini-Hochberg correction for the `Wald test`, `sGBJ`, `global boost test` and `global test`, respectively. Only the `global boost test` and `Wald` identified significant pathways among the 75 Grade 1 patients. No method identified significant pathways among the 119 Grade 3 patients, as shown in Panel B of Figure 3. Panel C of Figure 3 illustrates a general agreement between the methods, with the exception of the `global test`. As discussed in Section 4.1, asymptotic methods such as `sGBJ` and the `global test` can estimate p-values below 1e-3, a capability not achievable with the other methods using only 1,000 permutations.

## 5. Discussion

Gene set analysis methods for time-to-event data are relatively limited, and there is minimal guidance on selecting the most appropriate approach for specific contexts. To address this gap, we developed the `sGBJ` method, an extension of the `GBJ` test, tailored for gene set analysis in survival studies. To evaluate the effectiveness of `sGBJ` alongside other existing methods, we conducted a comprehensive benchmarking study aimed at identifying the most suitable approach under various conditions.

Our simulation studies revealed that the `sGBJ` method had a stringent control of type I error, which could reduce its statistical power, especially in scenarios involving a large number of genes. This issue was more pronounced compared to other methods, which generally demonstrated better power under similar conditions. Additionally, similar to Lee et al. (2011), we identified a key limitation of the `global test` method, which assumes that all gene effect sizes follow the same normal distribution – an assumption that may not always hold and could potentially limit the method’s effectiveness, as illustrated by our Type C scenario. In contrast, the `Wald test` consistently produced reliable results across different settings, while the `global boost test` excelled in scenarios where genes had large effect sizes (i.e., Type C). Importantly, all methods effectively controlled the type I error, underscoring the robustness of these approaches in maintaining statistical integrity.

Even though `sGBJ` demonstrates lower performance when the number of genes is high compared to the number of individuals, its power is comparable to other methods in most scenarios. Furthermore, in two real-world examples, this method identified a number of significant pathways on par with the other methods – while always maintaining type-I error control in simulation studies. Therefore, while `sGBJ` does not outperform previous methods, it appears as a sound and competitive alternative.

Our findings complement those reported by Lee et al. (2011), who compared the `global test`, `Wald test`, and `global boost test`. Their study found high power across all three methods; however, their simulation settings aligned with the `global test`’s assumption that all gene effects (i.e., *β*) are sampled from the same distribution. This likely contributed to the better performance of the `global test` in their simulations, particularly in contrast to our Type C scenario. In their real-world analysis of ovarian cancer data, Lee et al. (2011) observed that the `global test`, the `Wald test`, and the `global boost test` identified 12, 6, and 8 significant pathways, respectively. In our real-world data analysis, `Wald test`, `global boost test` and `sGBJ` consistently demonstrated high statistical power on both the Rembrandt and breast cancer datasets, while `global test` identified less significant genes.

Altogether, these findings suggest that both `Wald test` and `global boost test` should be prioritized as first-line approaches for pathway analysis. When the gene effects within a pathway are expected to be highly heterogeneous — resembling the Type C scenario and the breast cancer data — `global boost test` may offer slightly greater power, whereas `Wald test` could perform

better in more homogeneous settings. Meanwhile, `sGBJ` serves as a valuable complementary method, helping to validate and stabilize pathway discoveries.

The adaptation of `sGBJ` method to the time-to-event context is implemented in a `R` package available on the CRAN at https://CRAN.R-project.org/package=sGBJ. Code for this analysis is available at https://github.com/thomasferte/sGBJ_computation and on Zenodo (Ferté et al., 2025).

## Acknowledgements

Computer time for this study was provided by the computing facilities MCIA (Mésocentre de Calcul Intensif Aquitain) of the *Universé de Bordeaux* and of the *Université de Pau et des Pays de l’Adour*. This study was carried out in the framework of the University of Bordeaux’s France 2030 program / RRI PHDS.

## Data, script, code, and supplementary information availability

The adaptation of `sGBJ` method to the time-to-event context is implemented in a `R` package available on the CRAN at https://CRAN.R-project.org/package=sGBJ. Analysis code is available at https://github.com/thomasferte/sGBJ_computation and on Zenodo (Ferté et al., 2025). Data for Rembrandt study are available at https://www.ncbi.nlm.nih.gov/geo/query/acc.cgi?acc=GSE108474. Breast Cancer data are available from Markus Schroeder (2017).

## References

Adewale AJ, Dinu I, Potter JD, Liu Q, Yasui Y (2008). Pathway analysis of microarray data via regression. Journal of Computational Biology 15, 269–277. 10.1089/cmb.2008.0002.

Agniel D, Hejblum BP (2017). Variance component score test for time-course gene set analysis of longitudinal RNA-seq data. Biostatistics 18, 589–604. 10.1093/biostatistics/kxx005.

Ashburner M, Ball CA, Blake JA, Botstein D, Butler H, Cherry JM, Davis AP, Dolinski K, Dwight SS, Eppig JT, Harris MA, Hill DP, Issel-Tarver L, Kasarskis A, Lewis S, Matese JC, Richardson JE, Ringwald M, Rubin GM, Sherlock G (2000). Gene ontology: tool for the unification of biology. Nature genetics 25, 25–29. 10.1038/75556.

Barnett I, Mukherjee R, Lin X (2017). The generalized higher criticism for testing SNP-set effects in genetic association studies. Journal of the American Statistical Association 112, 64–76. 10.1080/01621459.2016.1192039.

Benjamini Y, Hochberg Y (1995). Controlling the false discovery rate: a practical and powerful approach to multiple testing. Journal of the Royal statistical society: series B (Methodological) 57, 289–300. 10.1111/j.2517-6161.1995.tb02031.x.

Benson JR, Jatoi I (2012). The global breast cancer burden. Future oncology 8, 697–702. 10.2217/fon.12.61.

Bolstad BM, Irizarry RA, Åstrand M, Speed TP (2003). A comparison of normalization methods for high density oligonucleotide array data based on variance and bias. Bioinformatics 19, 185–193. 10.1093/bioinformatics/19.2.185.

Boulesteix AL, Hothorn T (2010). Testing the additional predictive value of high-dimensional molecular data. BMC bioinformatics 11, 1–11. 10.1186/1471-2105-11-78.

Cai T, Tonini G, Lin X (2011). Kernel machine approach to testing the significance of multiple genetic markers for risk prediction. Biometrics 67, 975–986. 10.1111/j.1541-0420.2010.01544.x.

Chen R, Smith-Cohn M, Cohen AL, Colman H (2017). Glioma subclassifications and their clinical significance. Neurotherapeutics 14, 284–297. 10.1007/s13311-017-0519-x.

Dolgalev I (2025). msigdbr: MSigDB Gene Sets for Multiple Organisms in a Tidy Data Format. R package version 10.0.0. URL: https://igordot.github.io/msigdbr/.

Efron B, Tibshirani R (2007). On testing the significance of sets of genes. The annals of applied statistics 1, 107–129. 10.1214/07-AOAS101.

Engle RF (1984). Wald, likelihood ratio, and Lagrange multiplier tests in econometrics. Handbook of econometrics 2, 775–826. 10.1016/S1573-4412(84)02005-5.

Ferté T, Villain L, Thiébaut R, Hejblum BP (2025). thomasferte/sGBJ_computation: v1.1.0. 10.5281/zenodo.15196374. URL: https://zenodo.org/records/15196374 (visited on 04/11/2025).

Gao C, McDowell IC, Zhao S, Brown CD, Engelhardt BE (2016). Context Specific and Differential Gene Co-expression Networks via Bayesian Biclustering. PLOS Computational Biology 12. Ed. by Xianghong Jasmine Zhou, e1004791. 10.1371/journal.pcbi.1004791. URL: https://dx.plos.org/10.1371/journal.pcbi.1004791 (visited on 03/14/2025).

Gauthier M, Agniel D, Thiébaut R, Hejblum BP (2020). dearseq: a variance component score test for RNA-Seq differential analysis that effectively controls the false discovery rate. NAR Genomics and Bioinformatics 2, Lqaa093. 10.1093/nargab/lqaa093.

Gaynor SM, Sun R, Lin X, Quackenbush J (2019). Identification of differentially expressed gene sets using the Generalized Berk–Jones statistic. Bioinformatics 35, 4568–4576. 10.1093/bioinformatics/btz277.

Georgetown Database of Cancer (G-DOC) platform (n.d.). https://gdoc.georgetown.edu/gdoc/.

Goeman JJ, Oosting J, Cleton-Jansen AM, Anninga JK, Van Houwelingen HC (2005). Testing association of a pathway with survival using gene expression data. Bioinformatics 21, 1950–1957. 10.1093/bioinformatics/bti267.

Hejblum BP, Skinner J, Thiébaut R (2015). Time-course gene set analysis for longitudinal gene expression data. PLoS computational biology 11, e1004310. 10.1371/journal.pcbi.1004310.

Higham NJ (2002). Computing the nearest correlation matrix—a problem from finance. IMA Journal of Numerical Analysis 22, 329–343. 10.1093/imanum/22.3.329. eprint: https://academic.oup.com/imajna/article-pdf/22/3/329/2089524/220329.pdf. URL: https://doi.org/10.1093/imanum/22.3.329.

Ishwaran H, Kogalur UB, Blackstone EH, Lauer MS (2008). Random survival forests. The annals of applied statistics 2, 841–860. 10.1214/08-AOAS169.

Kanehisa M, Araki M, Goto S, Hattori M, Hirakawa M, Itoh M, Katayama T, Kawashima S, Okuda S, Tokimatsu T, Yamanishi Y (2007). KEGG for linking genomes to life and the environment. Nucleic acids research 36, D480–D484. 10.1093/nar/gkm882.

Law CW, Chen Y, Shi W, Smyth GK (2014). voom: Precision weights unlock linear model analysis tools for RNA-seq read counts. Genome biology 15, R29. 10.1186/gb-2014-15-2-r29.

Lee S, Kim J, Lee S (2011). A comparative study on gene-set analysis methods for assessing differential expression associated with the survival phenotype. BMC bioinformatics 12, 377. 10.1186/1471-2105-12-377.

Liberzon A, Subramanian A, Pinchback R, Thorvaldsdóttir H, Tamayo P, Mesirov JP (2011). Molecular signatures database (MSigDB) 3.0. Bioinformatics 27, 1739–1740. 10.1093/bioinformatics/btr260.

Love MI, Huber W, Anders S (2014). Moderated estimation of fold change and dispersion for RNAseq data with DESeq2. Genome biology 15, 550. 10.1186/s13059-014-0550-8.

Maciejewski H (2014). Gene set analysis methods: statistical models and methodological differences. Briefings in bioinformatics 15, 504–518. 10.1093/bib/bbt002.

Madhavan S, Gusev Y, Harris M, Tanenbaum DM, Gauba R, Bhuvaneshwar K, Shinohara A, Rosso K, Carabet LA, Song L, et al. (2011). G-DOC: a systems medicine platform for personalized oncology. Neoplasia 13, 771–783. 10.1593/neo.11806.

Madhavan S, Zenklusen JC, Kotliarov Y, Sahni H, Fine HA, Buetow K (2009). Rembrandt: helping personalized medicine become a reality through integrative translational research. Molecular cancer research 7, 157–167. 10.1158/1541-7786.mcr-08-0435.

Markus Schroeder BHK (2017). breastCancerNKI. 10.18129/B9.BIOC.BREASTCANCERNKI. URL: https://bioconductor.org/packages/breastCancerNKI (visited on 03/14/2025).

McPherson K, Steel C, Dixon J (2000). Breast cancer—epidemiology, risk factors, and genetics. Bmj 321, 624–628. 10.1136/bmj.321.7261.624.

Neykov M, Hejblum BP, Sinnott JA (2018). Kernel machine score test for pathway analysis in the presence of semi-competing risks. Statistical methods in medical research 27, 1099–1114. 10.1177/0962280216653427.

Norden AD, Wen PY (2006). Glioma therapy in adults. The neurologist 12, 279–292. 10.1097/01.nrl.0000250928.26044.47.

Ogata H, Goto S, Sato K, Fujibuchi W, Bono H, Kanehisa M (1999). KEGG: Kyoto encyclopedia of genes and genomes. Nucleic acids research 27, 29–34. 10.1093/nar/27.1.29.

Ostrom QT, Bauchet L, Davis FG, Deltour I, Fisher JL, Langer CE, Pekmezci M, Schwartzbaum JA, Turner MC, Walsh KM, Wrensch MR, Barnholtz-Sloan JS (2014). The epidemiology of glioma in adults: a “state of the science” review. Neuro-oncology 16, 896–913. 10.1093/neuonc/nou087.

Phipson B, Smyth G (2010). Permutation P-values Should Never Be Zero: Calculating Exact P-values When Permutations Are Randomly Drawn. Statistical Applications in Genetics and Molecular Biology 9. 10.2202/1544-6115.1585.

Robinson MD, McCarthy DJ, Smyth GK (2010). edgeR: a Bioconductor package for differential expression analysis of digital gene expression data. Bioinformatics 26, 139–140. 10.1093/bioinformatics/btp616.

Sathornsumetee S, Reardon DA, Desjardins A, Quinn JA, Vredenburgh JJ, Rich JN (2007). Molecularly targeted therapy for malignant glioma. Cancer 110, 13–24. 10.1002/cncr.22741.

Scully OJ, Bay BH, Yip G, Yu Y (2012). Breast cancer metastasis. Cancer Genomics-Proteomics 9, 311–320.

Subramanian A, Tamayo P, Mootha VK, Mukherjee S, Ebert BL, Gillette MA, Paulovich A, Pomeroy SL, Golub TR, Lander ES, Jill P. M (2005). Gene set enrichment analysis: a knowledge-based approach for interpreting genome-wide expression profiles. Proceedings of the National Academy of Sciences 102, 15545–15550. 10.1073/pnas.0506580102.

Sun R, Hui S, Bader GD, Lin X, Kraft P (2019). Powerful gene set analysis in GWAS with the Generalized Berk-Jones statistic. PLoS genetics 15, e1007530. 10.1371/journal.pgen.1007530.

Sun R, Lin X (2017). Set-based tests for genetic association using the generalized Berk-Jones statistic. arXiv preprint arXiv:1710.02469. 10.48550/arXiv.1710.02469.

Tibshirani R (1997). The lasso method for variable selection in the Cox model. Statistics in medicine 16, 385–395. 10.1002/(sici)1097-0258(19970228)16:4%3C385::aid-sim380%3E3.0.co;2-3.

Trevor Hastie RT (2017). impute. 10.18129/B9.BIOC.IMPUTE. URL: https://bioconductor.org/packages/impute (visited on 03/14/2025).

Van De Vijver MJ, He YD, Van’t Veer LJ, Dai H, Hart AA, Voskuil DW, Schreiber GJ, Peterse JL, Roberts C, Marton MJ, Parrish M, Atsma D, Witteveen A, Glas A, Delahaye L, Velde T, Bartelink H, Rodenhuis S, Rutgers ET, Friend SH, et al. (2002). A gene-expression signature as a predictor of survival in breast cancer. New England Journal of Medicine 347, 1999–2009. 10.1056/nejmoa021967.

Wang Z, Gerstein M, Snyder M (2009). RNA-Seq: a revolutionary tool for transcriptomics. Nature reviews genetics 10, 57–63. 10.1038/nrg2484.test

